# Filamentous bacteriophages induce proinflammatory responses in intestinal epithelial cells

**DOI:** 10.1101/2024.06.24.600496

**Authors:** Ambarish C. Varadan, Juris A. Grasis

## Abstract

Bacteriophages are the dominant members of the human enteric virome and can shape bacterial communities in the gut; however, our understanding of how they directly impact health and disease is limited. Previous studies have shown that specific bacteriophage populations are expanded in patients with Crohn’s disease (CD) and ulcerative colitis (UC), suggesting that fluctuations in the enteric virome may contribute to intestinal inflammation. Based on these studies, we hypothesized that a high bacteriophage burden directly induces intestinal epithelial responses. We found that filamentous inoviruses M13 and Fd induced dose-dependent IL-8 expression in the human intestinal epithelial cell line HT-29 to a greater degree than their lytic counterparts T4 and ϕX174 did. We also found that M13, but not Fd, reduced bacterial internalization in HT-29 cells. This led us to investigate the mechanism underlying M13-mediated inhibition of bacterial internalization by examining the antiviral and antimicrobial responses in these cells. M13 upregulated Type I and III IFN expression and augmented short-chain fatty acid (SCFA)-mediated LL-37 expression in HT-29 cells. Taken together, our data establish that inoviruses directly affect human intestinal epithelial cells. These results provide new insights into the complex interactions between bacteriophages and the intestinal mucosa, which may underlie disease pathogenesis.

## Introduction

The human enteric virome is composed of eukaryotic and prokaryotic viruses, including viruses that infect human cells, viruses that infect microbes (such as bacteria, fungi, and archaea), and plant viruses that are primarily derived from the environment and diet^1^. Alterations in the enteric virome have been reported in colorectal cancer^2–3^, inflammatory bowel disease^4–7^, obesity^8^, type I diabetes^9^, nonalcoholic fatty liver disease^10^, cystic fibrosis^11^, graft-versus-host disease^12^, as well as malnutrition^13^. Bacteriophages are the dominant component of the enteric virome^14^. These viruses infect bacteria and play a crucial role in shaping bacterial communities in mammalian systems^15^. Bacteriophages in healthy human intestines are predominantly temperate double-stranded DNA (dsDNA) *Caudovirales* or single-stranded DNA (ssDNA) *Microviridae*^16^. Metagenomic analysis reported that patients with Crohn’s disease (CD) and ulcerative colitis (UC) had a greater abundance of *Caudovirales* bacteriophages and fewer *Microviridae* bacteriophages within their intestines^4^, indicating that bacteriophage populations are altered in these disease states. However, the ability of bacteriophages to directly stimulate human intestinal epithelial cells has not been extensively explored. Although bacteriophages do not directly infect human cells, they do possess molecules that have been shown to stimulate the immune system^17^ and have been reported to elicit cytokines and antiviral responses in both murine and human leukocytes^18–20^. However, it remains unclear whether bacteriophages can directly stimulate the intestinal epithelium and potentially affect disease states in humans.

Gut bacteriophages consist of temperate phages located within bacterial genomes and free lytic bacteriophages associating with the intestinal mucus during the steady state^21^. Virulent bacteriophages follow a lytic lifecycle wherein each infection is followed by virion production and host cell lysis^21^. The lytic bacteriophages used in this study were T4 and ϕX174. T4 has been shown to induce the expansion of CD4+ and CD8+ T cells in Peyer’s patches of germ-free mice^19^. Previous studies have used bacteriophage ϕX174 as a T cell-dependent neoantigen for the assessment of antibody responses in patients^22–23^.

Temperate bacteriophages follow a lysogenic lifecycle wherein they integrate into the host bacterial chromosome as a prophage. Upon exposure to specific signals, including antibiotics^24^, short-chain fatty acids (SCFAs)^25^, reactive oxygen species^26^, temperature^27^, and compounds in foods^28^, prophages are induced, indicating that they re-enter the lytic cycle, causing bacterial lysis and phage release^21^. This emergent release of phages could potentially provide a source of antigenic stimuli for intestinal epithelial cells. However, a recent study reported the detection and characterization of novel inoviruses from gut commensal bacteria^29^. Filamentous bacteriophages (or inoviruses) are a subgroup of *Inoviridae*, a family of non-enveloped, single-stranded DNA bacteriophages. They infect both Gram-positive and Gram-negative bacterial species as well as some species of archaea^30^. A unique feature of filamentous bacteriophages is their ability to establish chronic lifecycles, wherein progeny virions are continuously extruded out of the bacterial cell envelope without lysing their host^31^. They adhere to either of two life cycles: episomally replicating phage or temperate phage that can integrate into the host chromosome^31^. The filamentous bacteriophages used in this study were M13 and Fd, which are episomally replicating phages. M13 has been shown to switch the immunosuppressive phenotype of tumor-associated macrophages (TAM) to an inflammatory M1 phenotype^32^. Bacteriophage Fd has been shown to stimulate TNF production in bone marrow-derived dendritic cells (BMDC)^18^. Temperate filamentous phages have previously been implicated in bacterial pathogenesis by contributing to biofilm formation^33^, increasing the virulence of wound infections, inhibiting cytokines important for bacterial clearance, and suppressing phagocytosis^18^. Despite the abundance and presence of bacteriophages at the mucosa, very few studies have been conducted on their direct impact on intestinal epithelial cells. In this study, we specifically investigated the mucosal and functional responses of intestinal epithelial HT-29 cells to lytic bacteriophages T4 and ϕX174, and filamentous bacteriophages M13 and Fd. Based on growing evidence for the ability of bacteriophages to interact with mammalian cells^18–20^, we hypothesized that an increased bacteriophage burden can stimulate mucosal responses in intestinal epithelial cells.

## Materials and Methods

### Bacterial strains and bacteriophage stocks

The bacterial strains and their respective bacteriophage stocks used in this study are listed in Table I. The four *E. coli* strains used in this study were obtained from the Leibniz Institute DSMZ-German Collection of Microorganisms and Cell Cultures GmbH (Leibniz, Germany). The strains were identified as follows: *Escherichia coli* B (DSM 613), *Escherichia coli* PC0886 (DSM 13127), *Escherichia coli* Lederberg (DSM 5695), and *Escherichia coli* LE392 (DSM 4230). Bacteria were cultured on 1% Luria Bertani (LB) (Fisher Scientific) agar (Fisher Scientific) plates and incubated overnight at 37 °C. One colony was subsequently used to inoculate a 50 ml tube containing 20 ml LB and incubated again overnight at 37 °C. The overnight culture was diluted with LB to an OD_600_ of 0.1 and then incubated at 37 °C for 2 h to reach an OD_600_ of 0.5 (corresponding to 10^6^ colony-forming units (CFU/mL)). This was determined to be the optimal bacterial titer for the double agar overlay plaque assay^34^ to propagate, as well as to determine the concentration of infectious bacteriophage particles.

**Table I.**
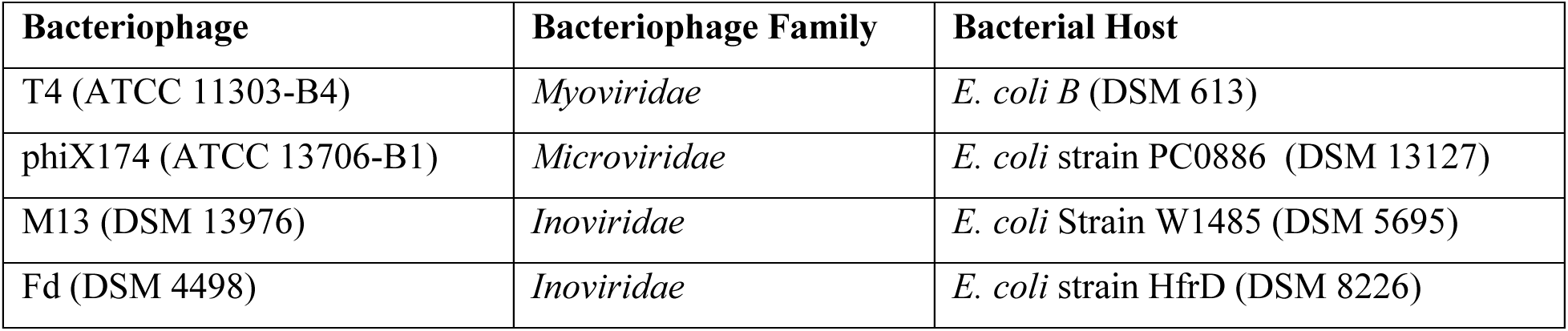
Bacteriophages and their respective bacterial strains used in this study. For each bacteriophage, the taxonomic family and bacterial host strains are presented.

### Bacteriophage purification

Endotoxin was removed following the Phage-on-Tap protocol^35^. Following filtration through a 100KDa Amicon filter unit (Fisher Scientific), the phage lysate was incubated with 1-Octanol at 4 °C for 2 h to remove endotoxins from the lysate. They were then dialyzed in 50-kDa Spectra Por Float-A-Lyzer G2 Dialysis Device (Cole-Parmer) against 15% Ethanol (24 hours) and 0.15 M NaCl (24 hours) prior to being quantified by double agar overlay plaque assays^34^. Bacteriophages M13 and Fd were harvested using polyethylene glycol-precipitation as described previously^36^. Bacteria were infected with stocks of bacteriophages at mid-log phase and cultured in 25 mL of LB broth for 16 h at 37 °C under shaking conditions. Bacteria were removed by centrifugation at 7,000 × *g* for 30 min, and the supernatant was treated with 1 μg/mL DNase I (Roche) for 2 h at 37 °C before 0.22 μm filtration. The virus-containing filtrate was then precipitated with 0.5 M NaCl and 8% PEG 8000 (Millipore Sigma) overnight at 4 °C. The phages were pelleted by centrifugation at 7,000× *g* for 30 min, and the pellet was suspended in sterile SM buffer. The suspension was centrifuged at 7,000 × *g* for 30 min, and the supernatant was subjected to another round of PEG precipitation. The purified filamentous phage pellets were suspended in sterile SM, incubated with 1-Octanol at 4 °C for 2 h to remove endotoxins from the precipitates, dialyzed in a 50-kDa Spectra Por Float-A-Lyzer G2 Dialysis Device (Cole-Parmer) against PBS for 48 h, and quantified using double agar overlay assays. Bacteriophage preparations were then tested for endotoxin by *Limulus* amoebocyte lysate (LAL) testing using the Pierce Chromogenic Endotoxin Quantification kit (Thermo Fisher Scientific).

### Bacteriophage Titer Determination

Bacteriophage titers were determined using the double agar overlay method^34^. Serial dilutions of the bacteriophage stocks were prepared. Phage dilutions (150 μL) were mixed with 200 μL of the respective bacterial host strain (10^8^ CFU/mL). Three milliliters of molten (45 °C) 0.5% LB Agar (Fisher Scientific) were added to the phage-bacteria mixture, plated onto Petri dishes, filled with a bottom layer of 1% LB agar, and incubated for 16 h at 37 °C. To determine the original bacteriophage concentration, plates containing one–100 distinguishable plaques were counted.

### Cell Culture

The human intestinal cell line HT-29 was obtained from the Cell Culture Facility at the University of California, Berkeley (Berkeley, CA, USA). HT-29 cells were cultured in Dulbecco’s modified Eagle’s medium (DMEM; Gibco; Thermo Fisher Scientific Inc.) supplemented with 10% (v/v) fetal bovine serum (FBS; Gibco; Thermo Fisher Scientific, Inc.), 100 U/mL Penicillin, and 100 µg/mL streptomycin (Gibco; Thermo Fisher Scientific, Inc.). Cells were incubated at 37 °C in a humidified atmosphere containing 5% CO2 (v/v). For activation and invasion assays, cells were seeded in 12-well tissue culture-treated plates (VWR) at a concentration of 2.0 x 10^5^ cells per well. Four days after seeding, the cells were serum starved for 24 h before conducting the experiments. For the kinetic assays, cells were treated with 10 ng/mL phorbol 12-myristate 13-acetate (Fisher Scientific) and 500 nM ionomycin (Fisher Scientific) for 3 h, after which they were washed (to remove any residual PMA and Ionomycin) and subsequently treated with either purified bacteriophage, LPS, a combination of bacteriophage and LPS, or SM buffer for specified time points. To evaluate antimicrobial peptide (AMP) expression, confluent HT-29 monolayers were treated with 0.5 mM Sodium Butyrate (Fisher Scientific).

### Reverse Transcription-quantitative Polymerase Chain Reaction (RT-qPCR)

Total RNA was extracted using TRIzol reagent (Thermo Fisher Scientific) according to the manufacturer’s protocol. cDNA was synthesized from 0.5 μg of total RNA by using Superscript III Reverse Transcriptase (Thermo Fisher Scientific). The primers used in this study are listed in Table III. Real-time RT-PCR was performed using a StepOnePlus thermocycler (Applied Biosystems). The final volume of the reaction cocktail was 20 μL, containing 1X PowerUP SYBR Green Master Mix (Thermo Fisher Scientific), 0.5 μM of each primer, and 1 μL of cDNA. The qRT-PCR protocol consisted of one step at 50 °C for 2 min (UDG activation) and 95 °C for 2 min (initial denaturation), followed by 40 cycles of amplification (95 °C for 15 s, 60 °C for 30 s, and 72 °C for 30 s). Data acquisition and analysis were performed using StepOne Plus Design and Analysis 2.0 software. The comparative 2^-ΔΔCt^ method was used to quantify gene expression level changes after normalization to the housekeeping gene GAPDH.

### Gentamicin Protection Assays

To determine whether inoviruses affected bacterial internalization by intestinal epithelial cells, gentamicin protection assays were performed as previously described^37^, with minor modifications. HT-29 cells were seeded in 12-well tissue culture-treated plates at a concentration of 3.0 x 10^5^ cells per well. Four days after seeding, the cells reached confluency and were serum-starved for 24 hours prior to conducting the experiments. Confluent monolayers were then washed three times with DMEM (without antibiotics or FBS) and stimulated with SM buffer, 1 × 10^10^ PFU/mL phage, 100 ng/mL LPS, or a combination of 1 × 10^10^ PFU/mL phage and 100 ng/mL LPS for 6 h. Cells were then washed with DMEM (without antibiotics or FBS) and challenged with overnight-diluted *E. coli* culture (10^7^ CFU/ml) at a multiplicity of infection (MOI) of 10:1 for 6 h. The cell monolayers were washed three times with DMEM (without antibiotics or FBS) prior to treatment with 100 µg/mL Gentamicin Sulfate (Fisher Scientific) for 1 h. The cells were then lysed using 0.1% Triton X-100 lysis buffer (Fisher Scientific). Serial dilutions of the lysates were plated on LB agar and bacterial colonies were counted after 16 h of incubation at 37 °C.

### Statistical Analyses

All experiments were conducted at least three times. Individual data points are displayed when possible and are represented as the mean ± Standard Error (SEM). Statistical significance was calculated using GraphPad PRISM software (version 10 for Windows; GraphPad Software, Inc.). Statistical significance was calculated using a two-tailed Student’s t-test, or ANOVA with Tukey’s or Dunnett’s multiple comparison correction, where two or more groups were compared. *P* < 0.05 was considered statistically significant.

## Results

### Bacteriophages activate proinflammatory cytokines in HT-29 epithelial cells

We assessed the immunogenicity of lytic bacteriophages T4 and ϕX174, and filamentous bacteriophages M13 and Fd in the colonic epithelial cell line HT-29, which is widely used to model the immune function of intestinal epithelial cells^38–39^. We hypothesized that bacteriophages induce an increase in the expression of the proinflammatory cytokine IL-8 in intestinal epithelial HT-29 cells. Our rationale for targeting IL-8 expression is that it is a major human chemokine that is rapidly induced in intestinal epithelial cells upon stimulation^40^. One challenge in studying mammalian immune responses to bacteriophages is the removal of endotoxins from the bacteriophage lysates. Bacterial endotoxins are highly immunogenic and can trigger inflammatory responses in TLR4-expressing mammalian cells^41^. We purified all bacteriophage lysates of endotoxin and determined their endotoxin concentrations to be less than 0.5 ng.ml^-1^ at the dilutions used in our experiments (Table II). Hence, exogenous LPS at a concentration of 0.5 ng.ml^-1^ was used as a control for our cellular activation experiments. Previous studies have shown that intestinal epithelial cells are hyporesponsive to LPS due to low surface expression of TLR4 and MD-2. In HT-29 cells, TLR4 protein is largely present in the cytoplasmic fraction, and the cells are hyporesponsive to LPS in an unprimed condition^42^. Therefore, we primed the cells with PMA/Ionomycin^43^ prior to bacteriophage treatment. To determine whether the observed immune response was induced by the bacteriophage rather than by possible endotoxin contamination present in the purified bacteriophage preparation, primed HT- 29 cells were stimulated with either 10^3^ PFU/HT-29 of bacteriophage, 0.5 ng.ml^-1^ LPS, a combination of purified bacteriophage (at a concentration of 10^3^ PFU/HT-29) and 0.5 ng.ml^-1^ exogenous LPS, or SM buffer for 2, 6, 12, and 24 h. As shown in Figure 1A, bacteriophage T4 induced significantly higher IL-8 expression compared to that induced by LPS, the combination of T4 and LPS, and SM buffer at 6 hours. No significant upregulation of IL-8 expression was observed in response to T4 treatment at any of the other time points. The addition of exogenous LPS to 10^3^ PFU/HT-29 bacteriophage T4 also did not lead to a significant difference in IL-8 expression induced by the phage alone at any of these time points. Interestingly, T4 reduced LPS-induced IL-8 activation at 6 h. Similar to T4, we observed the highest induction of IL-8 in response to stimulation with the bacteriophage ϕX174 at 6 h. We also observed that at 6 h, ϕX174 induced significantly higher IL-8 expression than that induced by the combination of ϕX174 and LPS (Figure 1B), indicating that lytic bacteriophages can counteract LPS-induced IL-8 expression at specific time points. To assess whether these bacteriophages could activate another proinflammatory cytokine, we evaluated TNFα expression at 6 h. Both T4 and ϕX174 induced significantly higher TNFα expression at 6 h compared to that induced by LPS and the combination of bacteriophage and LPS, although T4 induced greater TNFα expression compared to ϕX174 (Supplementary Figure 1A and 1B).

**Figure 1.**
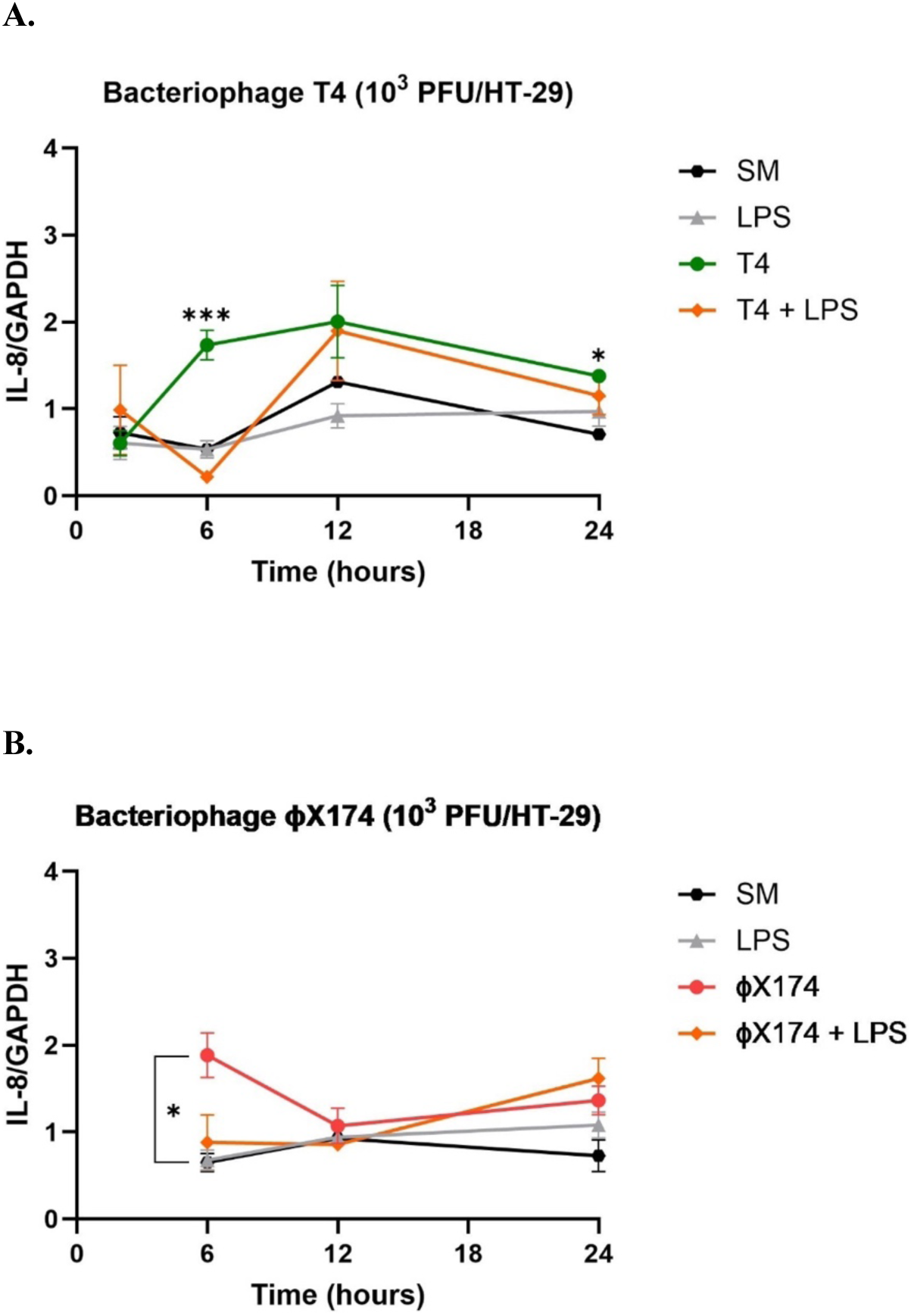
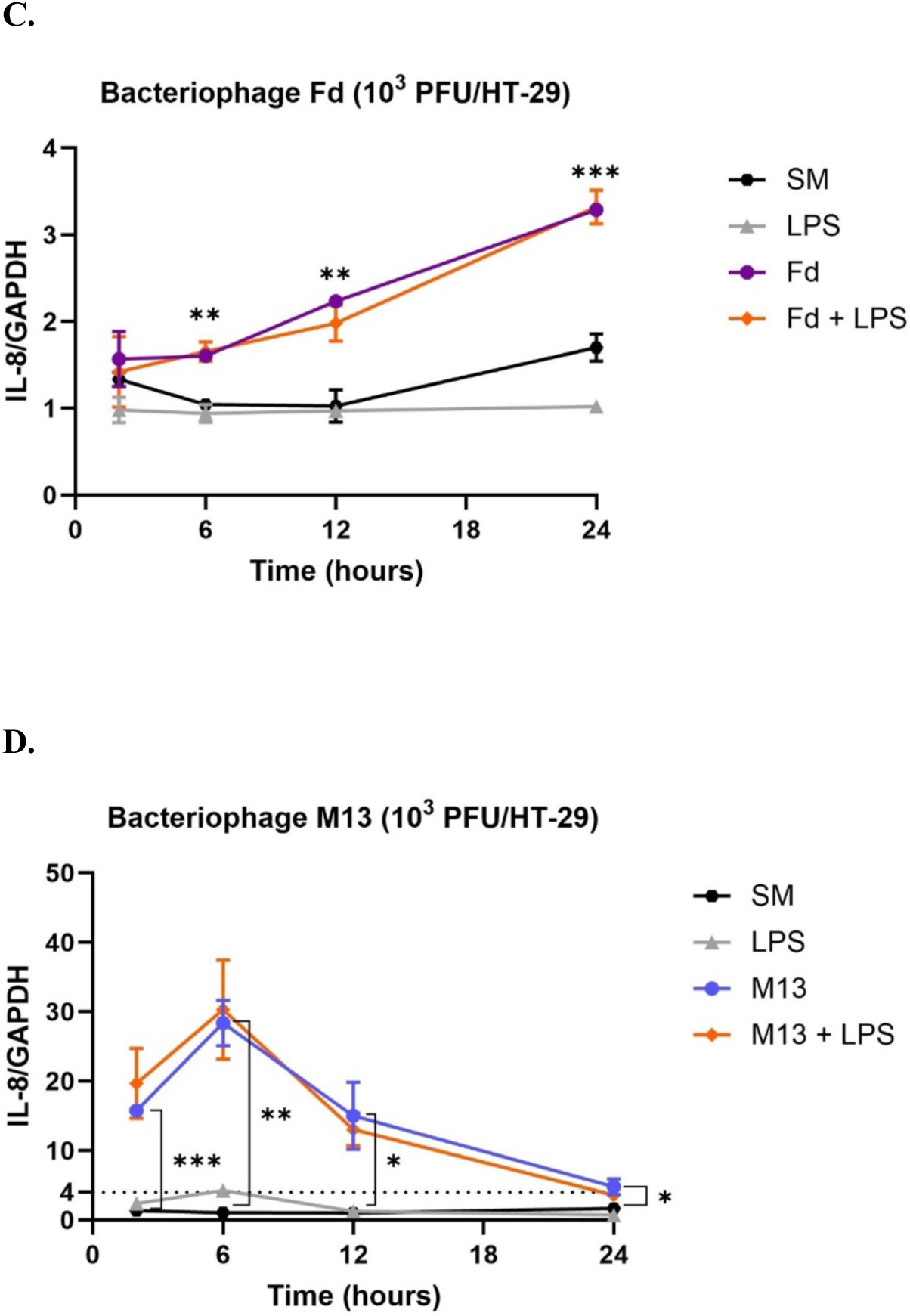
Kinetics of bacteriophage-mediated IL-8 activation of HT-29 epithelial cells. Primed HT-29 cells were stimulated with 10^9^ PFU/mL bacteriophages (A.) T4, (B.) ϕX174, (C.) Fd, and (D.) M13 for 2, 6, 12, and 24 hours. Respective controls for each experiment were SM buffer, LPS, bacteriophage, and a combination of bacteriophage with LPS. For each experiment, data was normalized to the expression level of GAPDH. All graphs are representative of n ≥ 3 experiments and depict the mean ± SEM of n ≥ 3 replicates. Analysis: one-way ANOVA with Tukey’s test for multiple comparisons. **\*** = P < 0.05, **\*\*** = P < 0.01, **\*\*\*** = P < 0.001, **ns** = not significant.

**Table II.** Titers of bacteriophage phage lysates before and after endotoxin removal.

| **Bacteriophage** | **Titer (PFU/mL)** |  | **Endotoxin (EU/mL)** |  |
| --- | --- | --- | --- | --- |
|  | **Lysate** | **Purified** | **Lysate** | **Purified** |
| T4 | 1.93 x 10<sup>11</sup> PFU/mL | 1.53 x 10<sup>10</sup> PFU/mL | 956.00 | 13.00 |
| phiX174 | 6.60 x 10<sup>10</sup> PFU/mL | 1.73 x 10<sup>10</sup> PFU/mL | 1508.58 | 16.34 |
| M13 | 1.50 x 10<sup>12</sup> PFU/mL | 3.17 x 10<sup>11</sup> PFU/mL | 3022.46 | 50.45 |
| Fd | 3.00 x 10<sup>13</sup> PFU/mL | 3.23 x 10<sup>11</sup> PFU/mL | 1565.20 | 71.03 |

**Table III.**
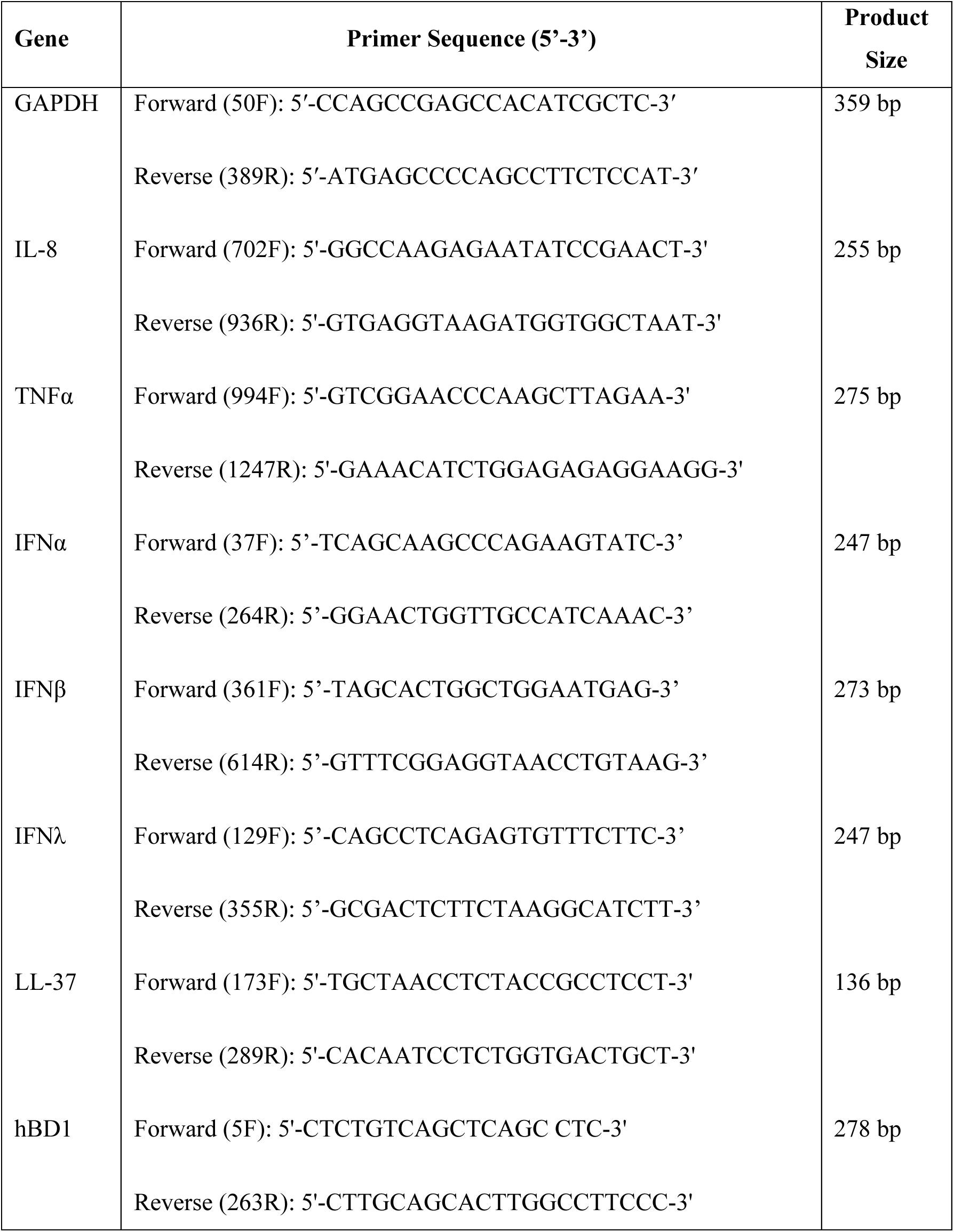
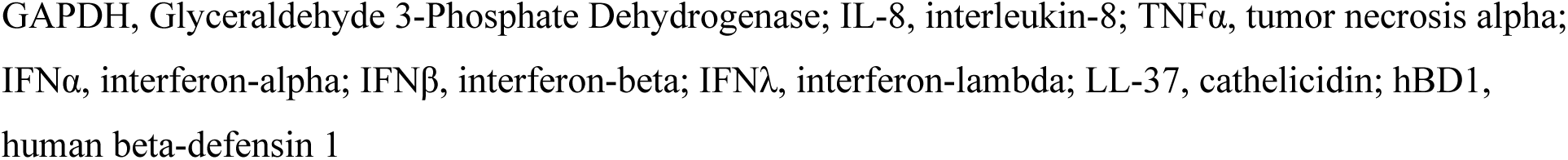
qRT-PCR primers used in this study.

We then evaluated the kinetics of IL-8 activation in HT-29 cells that were stimulated with either 10^3^ PFU/HT-29 of purified filamentous bacteriophage Fd, 0.5 ng/mL LPS, a combination of bacteriophage Fd (at a concentration of 10^3^ PFU/HT-29) and LPS, or SM buffer for 2, 6, 12, and 24 h (Figure 1C). At 6, 12, and 24 h post-treatment, bacteriophage Fd induced significantly higher IL-8 expression compared to that induced by LPS and SM buffer, with the highest IL-8 induction observed at 24 hours. The addition of exogenous LPS to bacteriophage Fd (at a concentration of 10^3^ PFU/HT-29) did not lead to a significant difference in IL-8 expression as induced by the phage alone at any of the time points. Next, we determined whether Fd-mediated IL-8 induction occurred in a concentration-dependent manner 24 h post-treatment (Supplementary Figure 2A). Lowering bacteriophage concentrations from 10^4^ PFU/HT-29 to 10^3^ PFU/HT-29 and from 10^3^ PFU/HT-29 to 10^2^ PFU/HT-29 cells significantly reduced IL-8 expression. However, no significant difference in IL-8 activation was observed in response to Fd at concentrations ranging from 10^2^-10^-1^ PFU/HT-29, demonstrating that Fd-mediated IL-8 activation occurred in a concentration-dependent manner (Supplementary Figure 2A).

The filamentous bacteriophage M13 induced significantly higher IL-8 expression compared to that induced by LPS and SM buffer at all evaluated time points, with the highest IL-8 induction observed at 6 h (Figure 1D). The addition of 0.5 ng/mL exogenous LPS to 10^3^ PFU/HT-29 of bacteriophage M13 did not lead to a significant difference in IL-8 expression induced by the phage alone at any of the time points. M13 also induced greater IL-8 activation than T4, ϕX174, or Fd. The dotted line in Figure 1D refers to the maximal IL-8 activation observed in response to other bacteriophages. Next, we determined whether M13-mediated IL-8 induction occurred in a concentration-dependent manner 24 h post-treatment (Supplementary Figure 2B). Lowering the bacteriophage M13 concentration from 10^4^ PFU/HT-29 to 10^2^ PFU/HT-29 did not significantly reduce IL-8 expression. Only when the bacteriophage concentration was reduced to 1 PFU/HT-29 and 0.1 PFU/HT-29 did we observe a significant reduction in IL-8 activation. In addition to IL-8, M13 induced significantly higher proinflammatory TNFα expression than that induced by LPS at 6 h post-treatment (Supplementary Figure 1C). The addition of exogenous LPS to bacteriophage M13 did not lead to a significant increase in TNFα expression. Collectively, these results suggest that bacteriophages can directly stimulate intestinal epithelial cells. Given that inoviruses induced a greater proinflammatory response compared to their lytic counterparts, we focus on intestinal epithelial responses to inoviruses M13 and Fd for the rest of this study.

### Inovirus M13 reduces bacterial internalization in gut epithelial cells

We next investigated whether bacteriophage M13 affects the bacterial infection rate through its effects on intestinal epithelial internalization. Bille *et al.* showed that the presence of filamentous bacteriophages results in increased bacterial colonization of epithelial cells^44^, implying that they can play pathogenic roles in bacterial infections of human cells. We hypothesized that the presence of filamentous bacteriophages would increase the number of bacteria that could be internalized by HT-29 cells. To test this, we incubated HT-29 cells with *E. coli* strain W1485 (host of M13) and M13 prior to measuring bacterial internalization. We found that the presence of M13 phage caused a significant reduction in the number of *E. coli* W1485 cells internalized by HT-29 cells (Supplementary Figure 3A). LPS has been shown to increase the permeability of intestinal epithelial tight junctions^45^. When HT-29 cells were co-stimulated with M13 and LPS prior to bacterial infection, we observed reduced bacterial internalization compared to HT-29 cells that were pre-stimulated with LPS prior to infection (Figure 2A). To determine whether M13 directly acts on HT-29 cells to inhibit bacterial internalization, we stimulated HT-29 cells with M13 prior to infecting them with *E. coli* 1485. We found that HT-29 cells stimulated with M13 internalized fewer *E. coli* 1485 cells than HT-29 cells stimulated with an equivalent volume of SM buffer (Figure 2A). Given that M13-mediated activation of HT-29 cells occurred in a concentration-dependent manner and was maintained over time (Supplementary Figure 2B), we hypothesized that M13-mediated inhibition of bacterial internalization would also occur in a concentration-dependent manner. However, we observed that stimulating HT-29 cells with different concentrations of M13 (10^0^-10^3^ PFU/HT-29) prior to bacterial infection did not significantly alter the number of internalized *E. coli* 1485 cells (Supplementary Figure 3B). However, when HT-29 cells were pre-stimulated with inovirus Fd prior to infection with *E. coli* strain HfrD (host of Fd), we did not observe a decrease in bacterial internalization compared to HT-29 cells that were pre-stimulated with an equivalent volume of SM buffer prior to infection (Figure 2B). Co-stimulation with Fd and LPS did not result in reduced internalization of *E. coli* HfrD compared to LPS stimulation alone. This demonstrates that the M13-mediated reduction in bacterial internalization was not universal across all inoviruses.

**Figure 2.**
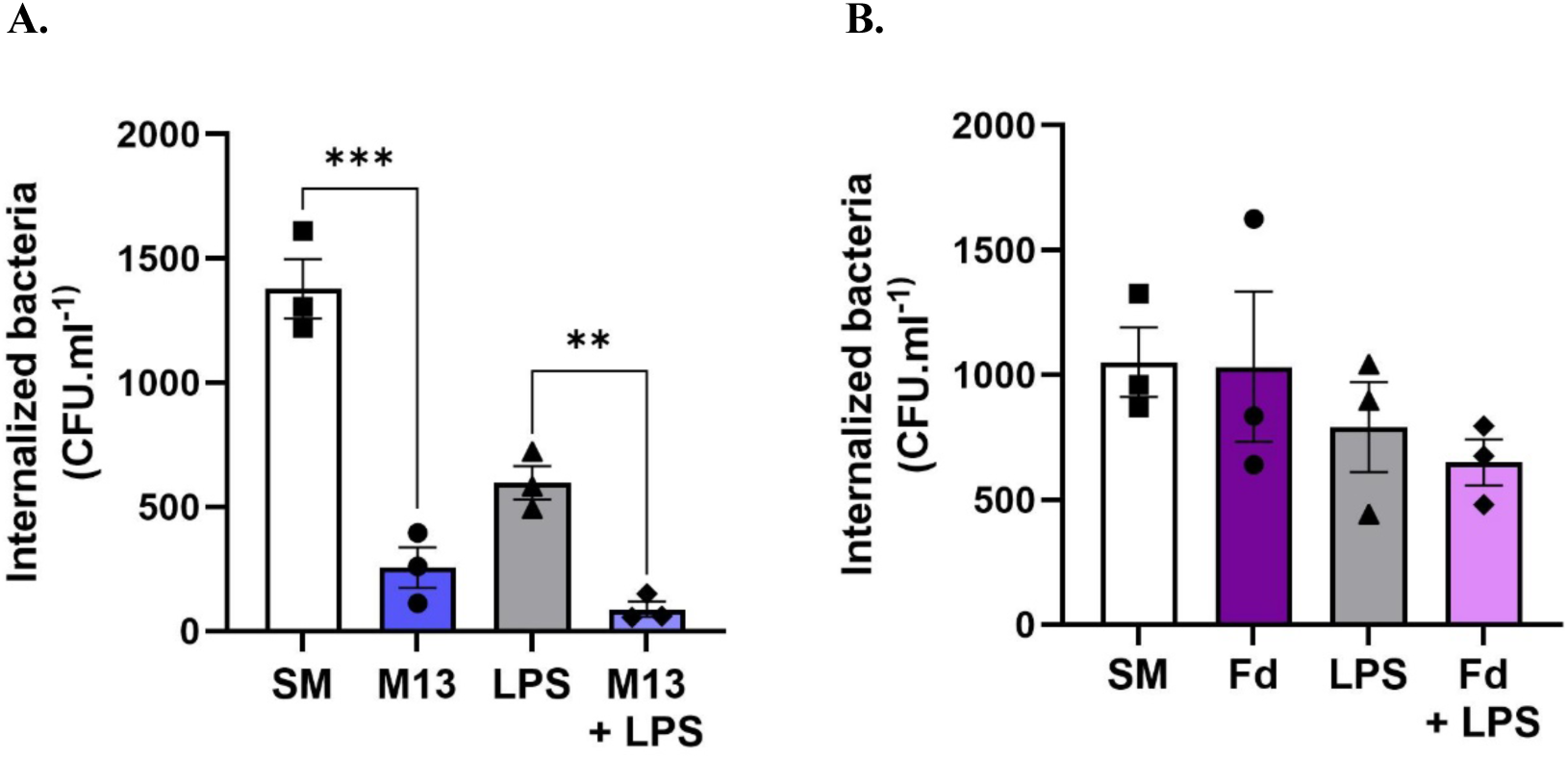
Inovirus M13 reduces bacterial internalization in gut epithelial cells. (A.) Confluent HT-29 monolayers were stimulated with SM buffer, M13 (10^4^ PFU/HT-29), LPS (100 ng/mL), and a combination of M13 (10^4^ PFU/HT-29) and LPS (100 ng/mL) for 6 hours prior to infection with *E. coli* W1485 (10^7^ PFU/mL). (B.) HT-29 cells were stimulated with SM buffer, Fd (10^4^ PFU/HT-29), LPS (100 ng/mL), and a combination of Fd (10^4^ PFU/HT-29) and LPS (100 ng/mL) for 6 hours prior to infection with *E. coli* HfrD (10^7^ PFU/mL). All graphs are representative of n ≥ 3 experiments and depict the mean ± SEM of n ≥ 3 replicates. Analysis: one-way ANOVA with Tukey’s test for multiple comparisons. **\*** = P < 0.05, **\*\*** = P < 0.01, **\*\*\*** = P < 0.001, **ns** = not significant.

### Bacteriophage M13 triggers antiviral Type I and Type III IFN responses

Next, we aimed to define the mechanism of the M13-mediated inhibition of bacterial internalization. Intestinal epithelial cells play a crucial role in maintaining intestinal homeostasis and regulating microbial colonization through a variety of mechanisms, including, antiviral^46^, antimicrobial^47^, and mucosal^48^ responses. Interferons (IFNs) are the main cytokines produced by intestinal cells, which control viral replication and spread within the body. The human intestinal epithelium exploits two types of IFNs for its protection: type I (IFN-α, IFN-β), and type III IFNs (IFN-λ1, -2, -3, and -4)^46^. However, Type I IFN signaling has also been shown to exert protective effects against bacterial infection^49–50^. Given that filamentous bacteriophages have been shown to promote the production of Type I Interferon (IFN) in murine BMDCs^18^, we hypothesized that they would induce Type I IFN responses in colonic epithelial HT-29 cells. As shown in Figure 3A, bacteriophage M13 induced significantly higher IFNβ expression than LPS, as well as the combination of LPS and M13 at 6 and 24 h. Bacteriophage M13 induced significantly higher IFNλ expression than LPS, as well as the combination of LPS and M13 at 6 h (Figure 3B). However, no significant change in IFNλ expression was observed in HT-29 cells stimulated with M13, LPS, or the combination of LPS and M13 at 24 h (Figure 3B). No significant change in IFNα expression was observed in HT-29 cells stimulated with M13, LPS, or the combination of LPS and M13 at 6 and 24 h (Figure 3C). No significant change in IFNβ, IFNλ, and IFNα expression was observed in HT-29 cells stimulated with either Fd, LPS, or the combination of LPS and Fd at either 6 or 24 hours (Figures 3D-F), demonstrating that inoviruses exert differential antiviral responses in intestinal epithelial cells.

**Figure 3.**
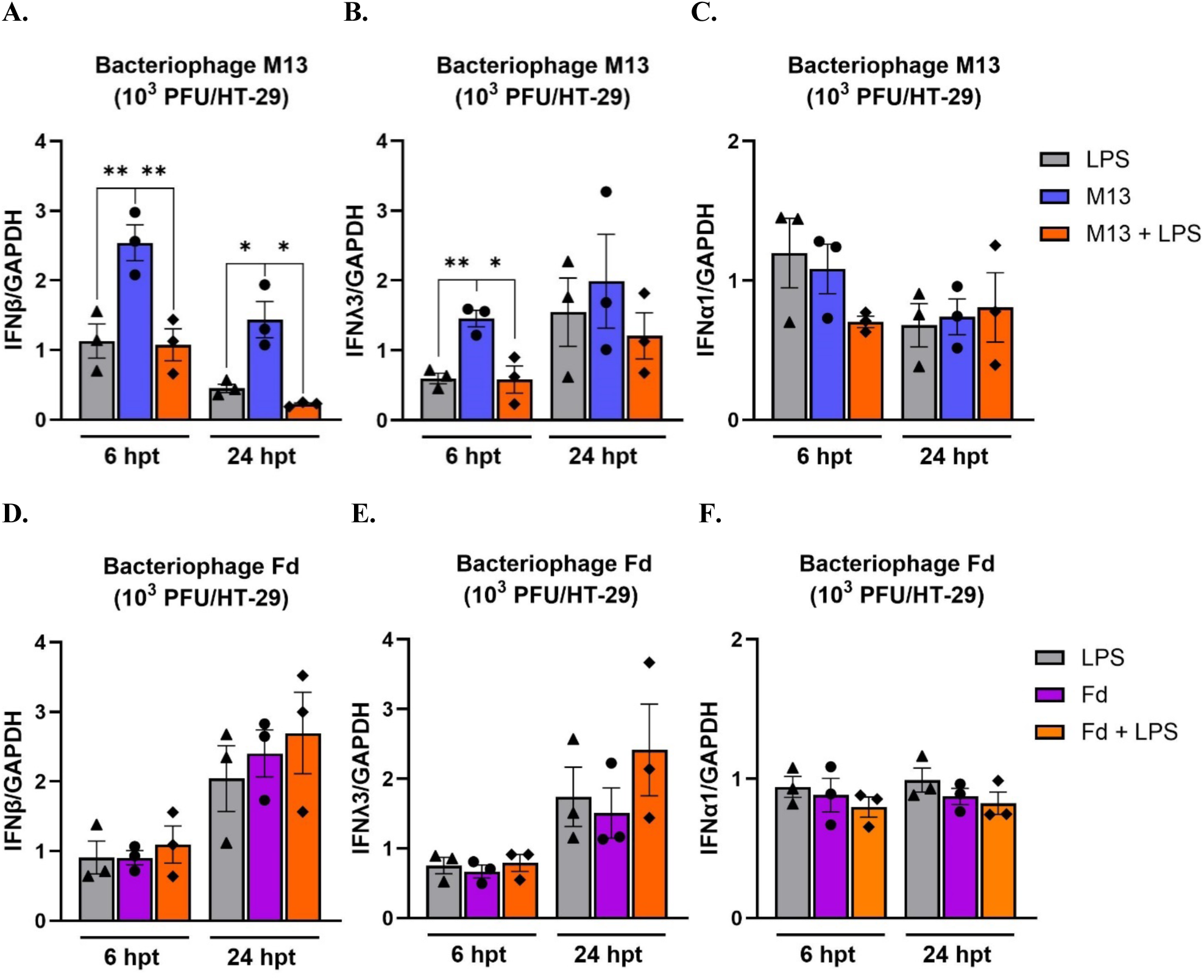
Bacteriophage M13 triggers antiviral Type I and Type III IFN responses. Effect of bacteriophage M13 stimulation on (A.) IFNβ, (B.) IFNλ, and (C.) IFNα induction at 6 and 24 hours, respectively. Effect of bacteriophage Fd stimulation on (D.) IFNβ, (E.) IFNλ, and (F.) IFNα induction at 6 and 24 hours, respectively. Respective controls for each experiment were LPS, bacteriophage, and a combination of bacteriophage with LPS. For each experiment, data was normalized to the expression level of GAPDH. All graphs are representative of n ≥ 3 experiments and depict the mean ± SEM of n ≥ 3 replicates. Analysis: one-way ANOVA with Tukey’s test for multiple comparisons. **\*** = P < 0.05, **\*\*** = P < 0.01, **\*\*\*** = P < 0.001, **ns** = not significant.

### Inovirus M13 augments butyrate-mediated LL-37 antimicrobial peptide expression

Based on a previous study that reported that pretreatment of mucus-producing intestinal epithelial cells with bacteriophages reduced subsequent bacterial attachment and cell death^51^, we hypothesized that inoviruses M13 and Fd induced antimicrobial peptide (AMP) gene expression in HT-29 cells.

Antimicrobial peptides (AMPs) are small (2-5 kDa), cationic, amphipathic peptides that play a critical role in innate immune defense mechanisms^52^ against a broad range of microorganisms, including bacteria, fungi, parasites, and viruses^47^. We determined whether bacteriophages induced the expression of cathelicidin LL-37 and β-defensin-1 (hβD-1) in HT-29 cells. LL-37 is a small, linear peptide that possesses broad bactericidal activity against both Gram-negative and Gram-positive bacteria^53^. Defensins are small, cationic peptides that contain disulfide bonds that are necessary to damage the bacterial cell membrane and eradicate bacteria^54^. β-Defensin hβD-1 is constitutively expressed in the gastrointestinal tract^55^. We found that neither M13 (Figure 4A-B) nor Fd (Figure 4C-D) significantly upregulated LL-37 or hβD-1 expression compared to buffer-treated HT-29 cells at 24 h post-treatment. Short-chain fatty acids (SCFAs) have been reported to be strong inducers of LL-37 expression in colonocytes^56–57^. They are microbial metabolites that constitute the major products of bacterial fermentation of dietary fiber in the intestines^58^. The major SCFAs produced in the colon are Acetate, Propionate, and Butyrate^59^. We then investigated whether inoviruses would affect AMP gene expression in the presence of butyrate. In agreement with previous studies^56–57^, the administration of butyrate alone significantly upregulated both LL-37 and hβD-1 expression in HT-29 cells. However, the administration of M13 along with butyrate induced a significantly higher LL-37 expression (Figure 4A), but not hβD-1 expression (Figure 4B), compared to that elicited by butyrate alone. The administration of exogenous LPS along with butyrate induced a similar expression of LL-37 as that induced by butyrate alone, confirming that the increased LL-37 expression in response to a combination of bacteriophage M13 and butyrate was not due to any residual LPS present in the bacteriophage preparation. The combination of Fd with butyrate did not significantly upregulate either LL-37 (Figure 4C) or hβD-1 expression (Figure 4D) compared to that elicited by butyrate alone. Collectively, these results suggest that specific bacteriophages can synergize with gut metabolites to induce AMP gene expression.

**Figure 4.**
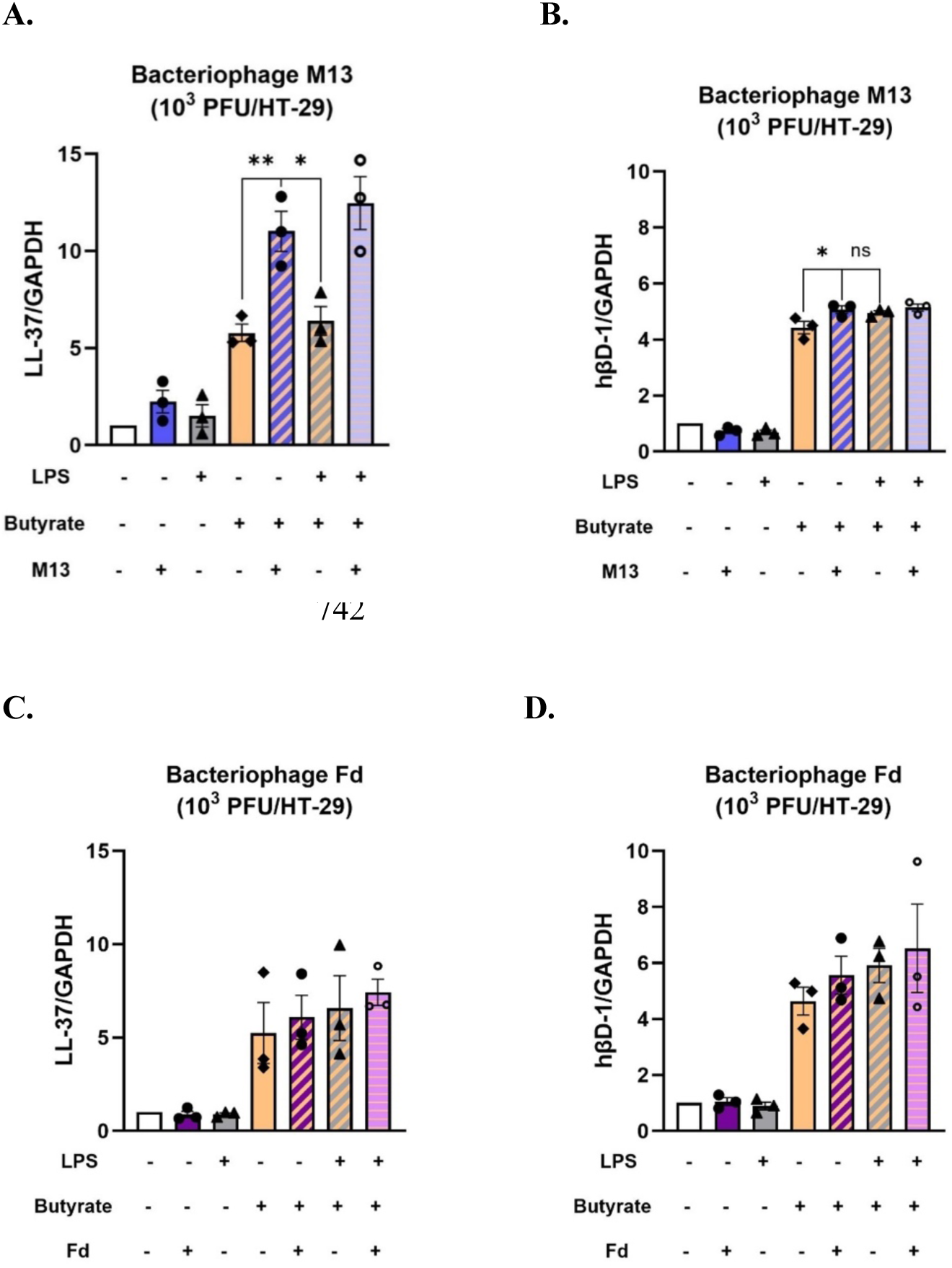
Inovirus M13 augments butyrate-mediated LL-37 antimicrobial peptide expression. LL-37 expression was assessed in response to bacteriophages (A.) M13 and (C.) Fd. hBD1 expression was evaluated in response to bacteriophages (B.) M13 and (D.) Fd. For each experiment, HT-29 cells were stimulated with SM buffer, inovirus, LPS, butyrate, or their combinations for 24 hours. Data was normalized to the expression level of GAPDH. All graphs are representative of n ≥ 3 experiments and depict mean with SEM of n ≥ 3 replicates. Analysis: one-way ANOVA with Tukey’s test for multiple comparisons. **\*** = P < 0.05, **\*\*** = P < 0.01, **\*\*\*** = P < 0.001, **ns** = not significant.

## DISCUSSION

Intestinal epithelial cells are the first line of defense and act as the interface between the intestinal microbiota and the body’s internal milieu^60^. Bacteriophages are the most abundant viruses in the gut microbiota and their abundance has been reported to be altered in many inflammatory disease states^2–3,8–13^ including IBD^4–7^. While previous studies have shown that bacteriophages can directly stimulate both human and murine phagocytes^18–20^, very few studies have been conducted to determine the direct impact of bacteriophages on human intestinal epithelial cells, which provide the frontline responses to gut microbiota for maintaining intestinal homeostasis. Understanding whether and how bacteriophages evoke intestinal epithelial cellular responses may be the first step in elucidating their roles in disease pathogenesis. In this study, we established that bacteriophages can directly stimulate proinflammatory responses in intestinal epithelial cells. These responses were dose-dependent, as in the case of inovirus M13 and Fd-mediated IL-8 activation, and greater than those induced by their lytic counterparts T4 and ϕX174. Previous studies have reported that bacteriophages can elicit both proinflammatory^61^ and anti- inflammatory responses^20^ in human cells, implying that their dynamics within the human host are both phage- and cell-specific. We observed differential dynamics between lytic bacteriophages and inoviruses regarding their synergy with LPS. Lytic bacteriophages (T4 and ϕX174), but not inoviruses (M13 and Fd), decreased the inflammatory response to LPS at 6 h post-treatment, as assessed by decreased intestinal epithelial expression of IL-8 and TNFα. Miernikiewicz *et al* previously reported that T4 short-tail fiber adhesin gp12 decreased LPS-induced proinflammatory cytokines IL-1α and IL-6 *in vivo*^62^. Furthermore, Zhang *et al.* reported that *Staphylococcus aureus* lytic bacteriophages suppressed LPS-induced inflammation in bovine mammary epithelial cells^63^. LPS is a component of the outer membrane of gram-negative *E. coli* and is one of the receptors for both bacteriophages T4^64^ and ϕX174^65^. The first step in phage infection is adsorption to the bacterial cell surface, which involves irreversible binding of T4 to LPS^64^. The mechanism(s) underlying bacteriophage modulation of LPS-induced proinflammatory immune responses needs to be investigated further and may provide new insights into the development of bacteriophages as a therapeutic option to combat antibiotic-resistant infections. We observed that inovirus M13 induced much higher IL-8 activation than the other phages. IL-8, a powerful chemoattractant released by IECs, attracts neutrophils to the basolateral surface of the epithelium^40^. Elevated IL-8 expression has been reported in the inflamed mucosa of patients with ulcerative colitis^66–68^. Determining the molecular pathways underlying inovirus-mediated IL-8 activation and whether this activation could increase neutrophil recruitment and inflammation will provide new insights into bacteriophage-induced immune responses and present an avenue for future research.

Previous studies have directly implicated inoviruses in the bacterial pathogenesis of human cells. Sweere *et al*^18^ reported that inovirus Pf4 impaired the clearance of *P. aeruginosa* by both murine and human phagocytes, whereas Bille *et al.* showed that the presence of inovirus MDAϕ resulted in increased colonization of *Neisseria meningitidis* on epithelial cells^44^. Here, we report that M13 reduces the internalization of *E. coli* W1485 in HT-29 cells. In contrast to the above-mentioned studies that used pathogenic bacteria such as *P. aeruginosa* and *N. meningitidis*, the bacterial strains of *E. coli* that we used for the internalization experiments were commensal. Common gut commensal species, such as *E. coli*^69^, can be internalized by enterocytes, although in significantly smaller numbers than invasive enteric pathogens (such as *Salmonella typhimurium* and *Listeria monocytogenes*)^70–72^. Here, we studied the invasiveness of *E. coli* W1485 by assessing its ability to be internalized by HT-29 cells. Although our data demonstrated that pre-stimulation of HT-29 cells with M13 reduced the internalization of *E. coli* W1485, we cannot conclude that M13 protects HT-29 cells against bacterial invasion. Internalization studies should be conducted with enteric pathogens to evaluate the protective potential of M13. We did not observe a significant reduction in bacterial internalization by HT-29 cells when they were pre-stimulated with inovirus Fd, suggesting that M13 and Fd may employ potentially differential interactions with colonic epithelial cells. The reason(s) behind this differential interaction is unclear and needs to be investigated further, but may be related to differences in their respective capsid protein structure or amino acid composition.

We sought to define the mechanism of the M13-mediated inhibition of bacterial internalization. M13 induced antiviral Type I IFN expression in HT-29 cells, corroborating previous studies demonstrating that inoviruses can trigger Type I IFN production in phagocytes^18^. Recently, single-cell RNA sequencing revealed that inoviruses upregulate several antiviral response genes in human basal epithelial cells (BCs), including IRF7^73^, a key transcription factor downstream of TLR3/TRIF signaling primarily induced by Type I IFNs and viral sensing^74^. Given that Type I IFN signaling has also been shown to have protective effects against bacterial infection^49–50^, it is likely that M13-mediated Type I IFN induction could play a role in regulating bacterial internalization by HT-29 cells. Alternatively, Sweere *et al.* demonstrated that inovirus Pf4 stimulated TLR3- and TRIF-dependent type I IFN production, inhibited TNF production, and limited phagocyte-mediated clearance of *P. aeruginosa*^18^. This is an example of an inovirus-mediated maladaptive antiviral response that results in impaired bacterial clearance and an increased establishment of infection. Conducting a global transcriptomic analysis of inovirus-stimulated intestinal epithelial cells will be informative, not only in determining which antiviral response genes are being induced, but also in providing insight(s) into the sensing mechanisms used by these cells to recognize inoviruses. This could be followed by functional studies to evaluate whether M13 could reduce bacterial internalization in intestinal epithelial cells lacking key components of the antiviral induction pathway. This would confirm whether M13-induced antiviral responses are protective or pathogenic in the context of bacterial infections of intestinal epithelial cells.

The key to delineating the mechanism by which M13 inhibits bacterial internalization by colonic epithelial cells may lie in the components of the mucosal surface. MUC2 is the main macromolecular component of intestinal mucus^75^, and previous reports have demonstrated that bacteriophages can adhere to^51^ and persist within mucosal surfaces^76^. This suggests that M13 interacts with and infects its host bacteria at the mucosal surface, thereby regulating bacterial internalization by intestinal epithelial cells through a mucus-dependent mechanism. Alternatively, Tian *et al.* reported that M13 can enter epithelial cells through clathrin-mediated endocytosis and macropinocytosis^77^. Given that both phages^18,77^ and commensal bacteria have been shown to be internalized by human cells, it is also likely that internalized M13 could regulate bacterial populations intracellularly^78^ through a mucus-independent mechanism.

In conclusion, these studies established that bacteriophages have direct effects on human intestinal epithelial cells and suggested that bacteriophages may play crucial roles in bacterial infections by directly interacting with intestinal epithelial cells.

## Disclosure statement

The authors declare no conflicts of interest in the generation, analysis, or reporting of this research.

## Data availability

The authors confirm that the data supporting the findings of this study are available within the article [and/or] its supplementary materials.

## Acknowledgments

We thank the members of the Grasis laboratory for related discussions on the generation of this manuscript. We would also like to thank Dr. David Gravano, technical director of the Stem Cell Instrumentation Foundry and Cytometry Center at UC Merced (SCIF).

## Funding source

This research was supported by the National Science Foundation (NSF) Biological Integration Institutes (BII): Host Virus Evolutionary Dynamics Institute (HVEDI; JAG 2119968).

---

**Supplementary Figure 1.**
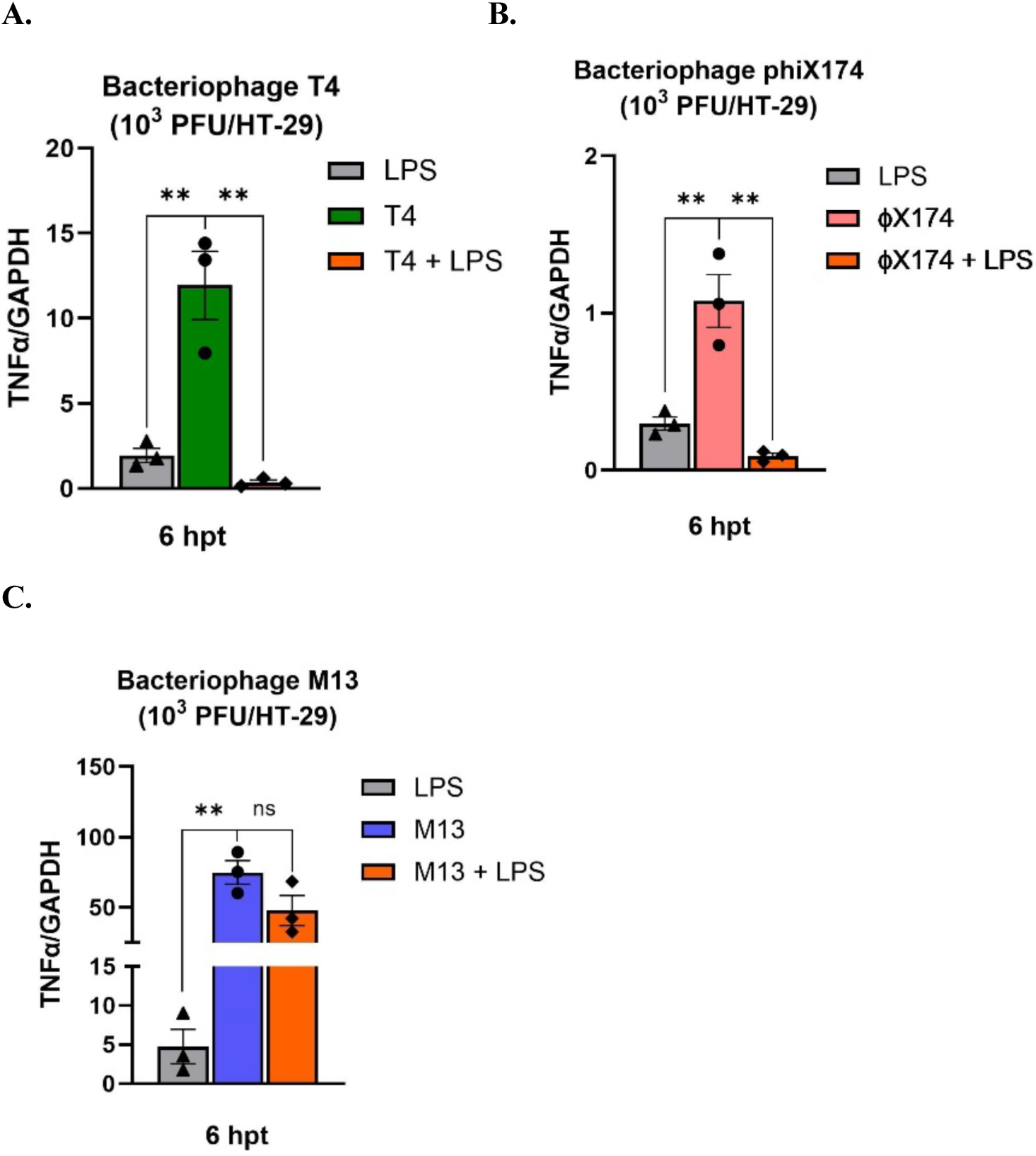
Bacteriophage-mediated induction of TNFα expression in HT-29 epithelial cells. TNFα expression was determined in primed HT-29 cells stimulated with 10^3^ PFU/HT-29 bacteriophage (A.) T4, (B.) ϕX174, and (C.) M13 for 6 hours. Respective controls for each experiment were LPS, bacteriophage, and a combination of bacteriophage with LPS. For each experiment, data was normalized to the expression level of GAPDH. All graphs are representative of n ≥ 3 experiments and depict mean with SEM of n ≥ 3 replicates. Analysis: one-way ANOVA with Tukey’s test for multiple comparisons. **\*** = P < 0.05, **\*\*** = P < 0.01, **\*\*\*** = P < 0.001, **ns** = not significant.

**Supplementary Figure 2.**
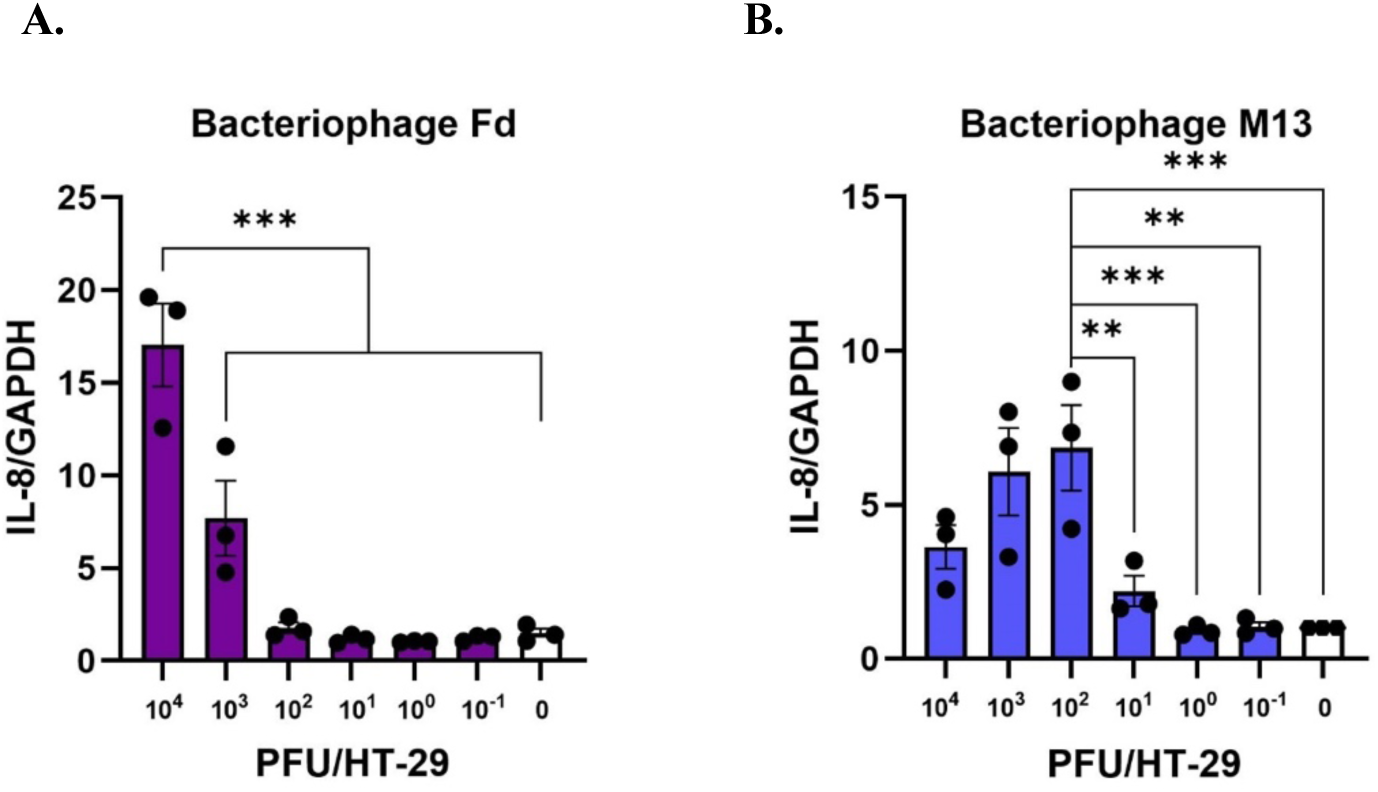
Inovirus dose-dependent IL-8 expression in HT-29 cells. in HT-29 cells in response to bacteriophages (A.) Fd and (B.) M13 at 24 hours. Concentrations were designated as a ratio of bacteriophages to HT-29 cells, and the ratios that were used ranged from 10^-1^-10^4^ bacteriophage PFU/HT-29. For each experiment, data was normalized to the expression level of GAPDH. All graphs are representative of n ≥ 3 experiments and depict mean with SEM of n ≥ 3 replicates. Analysis: one-way ANOVA with Dunnett’s test for multiple comparisons. **\*** = P < 0.05, **\*\*** = P < 0.01, **\*\*\*** = P < 0.001, **ns** = not significant.

**Supplementary Figure 3.**
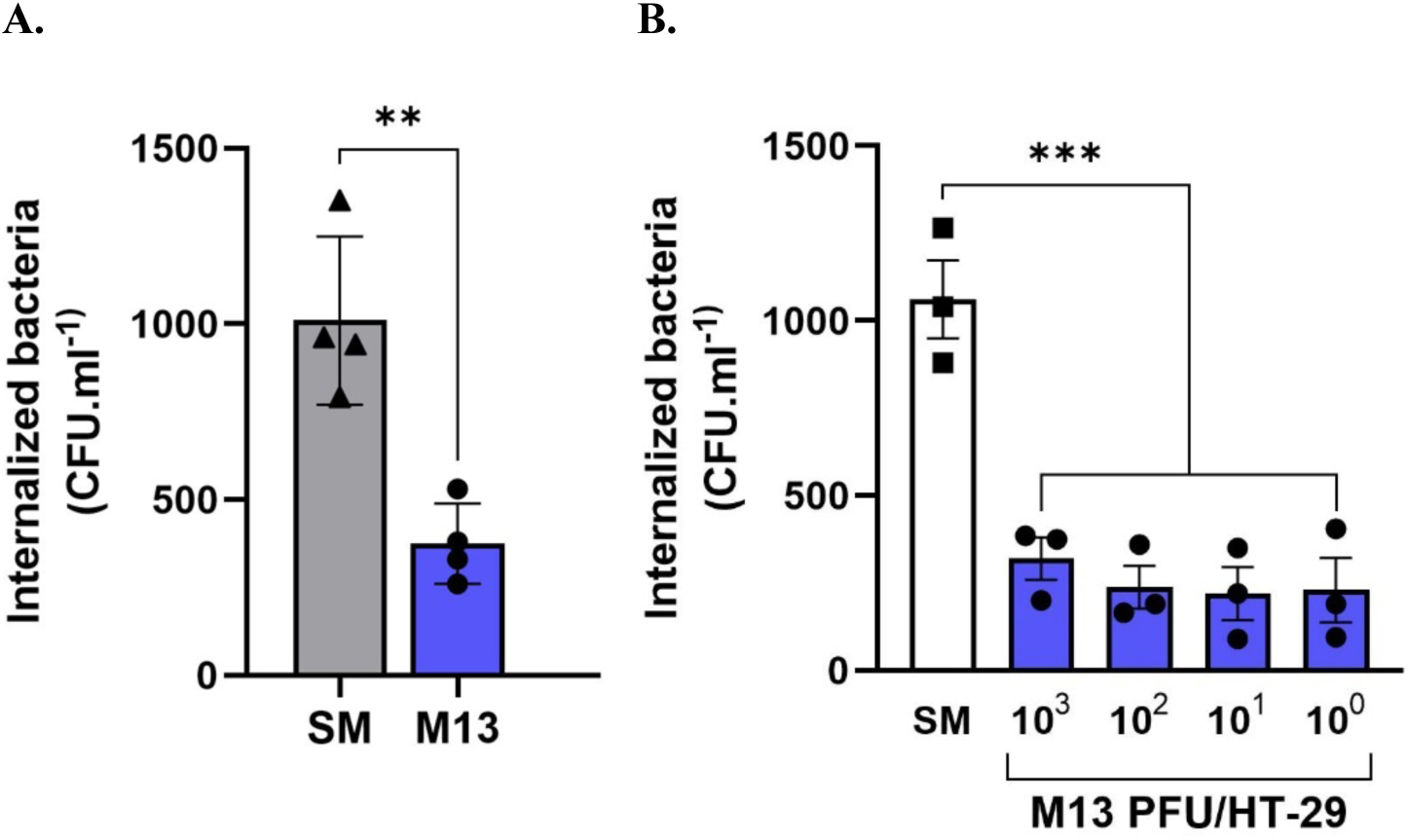
Bacteriophage M13 inhibition of epithelial bacterial internalization. (A.) Confluent HT-29 cells were incubated with 10^7^ CFU/mL *E. coli* strain W1485 and 10^7^ PFU/mL M13 (phage-bacteria ratio or MOI =1) for 6 h prior to determining bacterial internalization. (B.) HT-29 cells were stimulated with different concentrations of M13 (10^0^-10^3^ PFU/HT-29) for 6 h prior to infection with 10^7^ PFU/mL *E. coli* W1485 for 6 h. All graphs are representative of n ≥ 3 experiments and depict the mean ± SEM of n ≥ 3 replicates. Analysis: (A.) Two-tailed Student’s t test; (B.) one-way ANOVA with Tukey’s test for multiple comparisons. **\*** = P < 0.05, **\*\*** = P < 0.01, **\*\*\*** = P < 0.001, **ns** = not significant.

## Notes

### Competing Interest Statement

The authors have declared no competing interest.

## References

1. Liang, G., & Bushman, F. D. (2021). The human virome: Assembly, composition and host interactions. Nature Reviews Microbiology, 19(8), Article 8. 10.1038/s41579-021-00536-5

2. Hannigan, G. D., Duhaime, M. B., Ruffin, M. T., Koumpouras, C. C., & Schloss, P. D. (2018). Diagnostic Potential and Interactive Dynamics of the Colorectal Cancer Virome. mBio, 9(6), 10.1128/mbio.02248-18. 10.1128/mbio.02248-18

3. Nakatsu, G., Zhou, H., Wu, W. K. K., Wong, S. H., Coker, O. O., Dai, Z., Li, X., Szeto, C.-H., Sugimura, N., Lam, T. Y.-T., Yu, A. C.-S., Wang, X., Chen, Z., Wong, M. C.-S., Ng, S. C., Chan, M. T. V., Chan, P. K. S., Chan, F. K. L., Sung, J. J.-Y., & Yu, J. (2018). Alterations in Enteric Virome Are Associated With Colorectal Cancer and Survival Outcomes. Gastroenterology, 155(2), 529–541.e5. 10.1053/j.gastro.2018.04.018. .

4. Norman, J. M., Handley, S. A., Baldridge, M. T., Droit, L., Liu, C. Y., Keller, B. C., Kambal, A., Monaco, C. L., Zhao, G., Fleshner, P., Stappenbeck, T. S., McGovern, D. P. B., Keshavarzian, A., Mutlu, E. A., Sauk, J., Gevers, D., Xavier, R. J., Wang, D., Parkes, M., & Virgin, H. W. (2015). Disease-Specific Alterations in the Enteric Virome in Inflammatory Bowel Disease. Cell, 160(3), 447–460. 10.1016/j.cell.2015.01.002

5. Clooney, A. G., Sutton, T. D. S., Shkoporov, A. N., Holohan, R. K., Daly, K. M., O’Regan, O., Ryan, F. J., Draper, L. A., Plevy, S. E., Ross, R. P., & Hill, C. (2019). Whole-Virome Analysis Sheds Light on Viral Dark Matter in Inflammatory Bowel Disease. Cell Host & Microbe, 26(6), 764–778.e5. 10.1016/j.chom.2019.10.009

6. Duerkop, B. A., Kleiner, M., Paez-Espino, D., Zhu, W., Bushnell, B., Hassell, B., Winter, S. E., Kyrpides, N. C., & Hooper, L. V. (2018). Murine colitis reveals a disease-associated bacteriophage community. Nature Microbiology, 3(9), Article 9. 10.1038/s41564-018-0210-y.

7. Liang, G., Conrad, M. A., Kelsen, J. R., Kessler, L. R., Breton, J., Albenberg, L. G., Marakos, S., Galgano, A., Devas, N., Erlichman, J., Zhang, H., Mattei, L., Bittinger, K., Baldassano, R. N., & Bushman, F. D. (2020). Dynamics of the Stool Virome in Very Early-Onset Inflammatory Bowel Disease. Journal of Crohn’s and Colitis, 14(11), 1600–1610. 10.1093/ecco-jcc/jjaa094.

8. Yang, K., Niu, J., Zuo, T., Sun, Y., Xu, Z., Tang, W., Liu, Q., Zhang, J., Ng, E. K. W., Wong, S. K. H., Yeoh, Y. K., Chan, P. K. S., Chan, F. K. L., Miao, Y., & Ng, S. C. (2021). Alterations in the Gut Virome in Obesity and Type 2 Diabetes Mellitus. Gastroenterology, 161(4), 1257–1269.e13. 10.1053/j.gastro.2021.06.056

9. Zhao, G., Vatanen, T., Droit, L., Park, A., Kostic, A. D., Poon, T. W., Vlamakis, H., Siljander, H., Härkönen, T., Hämäläinen, A.-M., Peet, A., Tillmann, V., Ilonen, J., Wang, D., Knip, M., Xavier, R. J., & Virgin, H. W. (2017). Intestinal virome changes precede autoimmunity in type I diabetes-susceptible children. Proceedings of the National Academy of Sciences, 114(30), E6166–E6175. 10.1073/pnas.1706359114

10. Lang, S., Demir, M., Martin, A., Jiang, L., Zhang, X., Duan, Y., Gao, B., Wisplinghoff, H., Kasper, P., Roderburg, C., Tacke, F., Steffen, H.-M., Goeser, T., Abraldes, J. G., Tu, X. M., Loomba, R., Stärkel, P., Pride, D., Fouts, D. E., & Schnabl, B. (2020). Intestinal Virome Signature Associated With Severity of Nonalcoholic Fatty Liver Disease. Gastroenterology, 159(5), 1839–1852. 10.1053/j.gastro.2020.07.005. .

11. Willner, D., Furlan, M., Haynes, M., Schmieder, R., Angly, F. E., Silva, J., Tammadoni, S., Nosrat, B., Conrad, D., & Rohwer, F. (2009). Metagenomic Analysis of Respiratory Tract DNA Viral Communities in Cystic Fibrosis and Non-Cystic Fibrosis Individuals. PLOS ONE, 4(10), e7370. 10.1371/journal.pone.0007370. .

12. Legoff, J., Resche-Rigon, M., Bouquet, J., Robin, M., Naccache, S. N., Mercier-Delarue, S., Federman, S., Samayoa, E., Rousseau, C., Piron, P., Kapel, N., Simon, F., Socié, G., & Chiu, C. Y. (2017). The eukaryotic gut virome in hematopoietic stem cell transplantation: New clues in enteric graft-versus-host disease. Nature Medicine, 23(9), Article 9. 10.1038/nm.4380. .

13. Khan Mirzaei, M., Khan, Md. A. A., Ghosh, P., Taranu, Z. E., Taguer, M., Ru, J., Chowdhury, R., Kabir, Md. M., Deng, L., Mondal, D., & Maurice, C. F. (2020). Bacteriophages Isolated from Stunted Children Can Regulate Gut Bacterial Communities in an Age-Specific Manner. Cell Host & Microbe, 27(2), 199–212.e5. 10.1016/j.chom.2020.01.004

14. Hoyles, L., McCartney, A. L., Neve, H., Gibson, G. R., Sanderson, J. D., Heller, K. J., & van Sinderen, D. (2014). Characterization of virus-like particles associated with the human fecal and cecal microbiota. Research in Microbiology, 165(10), 803–812. 10.1016/j.resmic.2014.10.006.

15. Hsu, B. B., Gibson, T. E., Yeliseyev, V., Liu, Q., Lyon, L., Bry, L., Silver, P. A., & Gerber, G. K. (2019). Dynamic Modulation of the Gut Microbiota and Metabolome by Bacteriophages in a Mouse Model. Cell Host & Microbe, 25(6), 803–814.e5. 10.1016/j.chom.2019.05.001.

16. Minot, S., Sinha, R., Chen, J., Li, H., Keilbaugh, S. A., Wu, G. D., Lewis, J. D., & Bushman, F. D. (2011). The human gut virome: Inter-individual variation and dynamic response to diet. Genome Research, 21(10), 1616–1625. 10.1101/gr.122705.111.

17. Duerkop, B. A., & Hooper, L. V. (2013). Resident viruses and their interactions with the immune system. Nature Immunology, 14(7), Article 7. 10.1038/ni.2614.

18. Sweere JM, Van Belleghem JD, Ishak H, Bach MS, Popescu M, Sunkari V, Kaber G, Manasherob R, Suh GA, Cao X, de Vries CR, Lam DN, Marshall PL, Birukova M, Katznelson E, Lazzareschi DV, Balaji S, Keswani SG, Hawn TR, Secor PR, Bollyky PL. (2019). Bacteriophage trigger antiviral immunity and prevent clearance of bacterial infection. 363(6434):eaat9691. doi: 10.1126/science.aat9691. PMID: 30923196; PMCID: PMC6656896.

19. Gogokhia, L., Buhrke, K., Bell, R., Hoffman, B., Brown, D. G., Hanke-Gogokhia, C., Ajami, N. J., Wong, M. C., Ghazaryan, A., Valentine, J. F., Porter, N., Martens, E., O’Connell, R., Jacob, V., Scherl, E., Crawford, C., Stephens, W. Z., Casjens, S. R., Longman, R. S., & Round, J. L. (2019). Expansion of Bacteriophages Is Linked to Aggravated Intestinal Inflammation and Colitis. Cell Host & Microbe, 25(2), 285–299.e8. 10.1016/j.chom.2019.01.008

20. Van Belleghem, J. D., Clement, F., Merabishvili, M., Lavigne, R., & Vaneechoutte, M. (2017). Pro- and anti-inflammatory responses of peripheral blood mononuclear cells induced by Staphylococcus aureus and Pseudomonas aeruginosa phages. Scientific Reports, 7(1), Article 1. 10.1038/s41598-017-08336-9.

21. Sausset, R., Petit, M. A., Gaboriau-Routhiau, V., & De Paepe, M. (2020). New insights into intestinal phages. Mucosal Immunology, 13(2), 205–215. 10.1038/s41385-019-0250-5.

22. Pyun, K. H., Ochs, H. D., Wedgwood, R. J., Xiqiang, Y., Heller, S. R., & Reimer, C. B. (1989). Human antibody responses to bacteriophage φX 174: Sequential induction of IgM and IgG subclass antibody. Clinical Immunology and Immunopathology, 51(2), 252–263. 10.1016/0090-1229(89)90024-X

23. Pescovitz, M. D., Torgerson, T. R., Ochs, H. D., Ocheltree, E., McGee, P., Krause-Steinrauf, H., Lachin, J. M., Canniff, J., Greenbaum, C., Herold, K. C., Skyler, J. S., Weinberg, A., & Type 1 Diabetes TrialNet Study Group. (2011). Effect of rituximab on human in vivo antibody immune responses. The Journal of Allergy and Clinical Immunology, *128*(6), 1295-1302.e5. 10.1016/j.jaci.2011.08.008

24. Sutcliffe, S. G., Shamash, M., Hynes, A. P., & Maurice, C. F. (2021). Common Oral Medications Lead to Prophage Induction in Bacterial Isolates from the Human Gut. Viruses, 13(3), 455. 10.3390/v13030455

25. Oh, J. H., Alexander, L. M., Pan, M., Schueler, K. L., Keller, M. P., Attie, A. D., Walter, J., & van Pijkeren, J. P. (2019). Dietary Fructose and Microbiota-Derived Short-Chain Fatty Acids Promote Bacteriophage Production in the Gut Symbiont Lactobacillus reuteri. Cell host & microbe, 25(2), 273–284.e6. 10.1016/j.chom.2018.11.016

26. Diard, M., Bakkeren, E., Cornuault, J. K., Moor, K., Hausmann, A., Sellin, M. E., Loverdo, C., Aertsen, A., Ackermann, M., De Paepe, M., Slack, E., & Hardt, W. D. (2017). Inflammation boosts bacteriophage transfer between *Salmonella* spp. *Science (New York*, N.Y.), 355(6330), 1211–1215. 10.1126/science.aaf8451

27. Henrot, C., & Petit, M. A. (2022). Signals triggering prophage induction in the gut microbiota. Molecular microbiology, 118(5), 494–502. 10.1111/mmi.14983

28. Boling, L., Cuevas, D. A., Grasis, J. A., Kang, H. S., Knowles, B., Levi, K., Maughan, H., McNair, K., Rojas, M. I., Sanchez, S. E., Smurthwaite, C., & Rohwer, F. (2020). Dietary prophage inducers and antimicrobials: toward landscaping the human gut microbiome. Gut microbes, 11(4), 721–734. 10.1080/19490976.2019.1701353

29. Burckhardt, J. C., Chong, D. H. Y., Pett, N., & Tropini, C. (2023). Gut commensal Enterocloster species host inoviruses that are secreted in vitro and in vivo. Microbiome, 11(1), 65. 10.1186/s40168-023-01496-z.

30. Roux, S., Krupovic, M., Daly, R. A., Borges, A. L., Nayfach, S., Schulz, F., Sharrar, A., Matheus Carnevali, P. B., Cheng, J.-F., Ivanova, N. N., Bondy-Denomy, J., Wrighton, K. C., Woyke, T., Visel, A., Kyrpides, N. C., & Eloe-Fadrosh, E. A. (2019). Cryptic inoviruses revealed as pervasive in bacteria and archaea across Earth’s biomes. Nature Microbiology, 4(11), Article 11. 10.1038/s41564-019-0510-x

31. Rakonjac, J., Bennett, N. J., Spagnuolo, J., Gagic, D., & Russel, M. (2011). Filamentous Bacteriophage: Biology, Phage Display and Nanotechnology Applications. Current Issues in Molecular Biology, 13(2), Article 2. 10.21775/cimb.013.051

32. Eriksson, F., Tsagozis, P., Lundberg, K., Parsa, R., Mangsbo, S. M., Persson, M. A. A., Harris, R. A., & Pisa, P. (2009). Tumor-Specific Bacteriophages Induce Tumor Destruction through Activation of Tumor-Associated Macrophages1. The Journal of Immunology, 182(5), 3105–3111. 10.4049/jimmunol.0800224.

33. Secor, P. R., Sweere, J. M., Michaels, L. A., Malkovskiy, A. V., Lazzareschi, D., Katznelson, E., Rajadas, J., Birnbaum, M. E., Arrigoni, A., Braun, K. R., Evanko, S. P., Stevens, D. A., Kaminsky, W., Singh, P. K., Parks, W. C., & Bollyky, P. L. (2015). Filamentous Bacteriophage Promote Biofilm Assembly and Function. Cell Host & Microbe, 18(5), 549–559. 10.1016/j.chom.2015.10.013.

34. Kropinski, A. M., Mazzocco, A., Waddell, T. E., Lingohr, E., & Johnson, R. P. (2009). Enumeration of Bacteriophages by Double Agar Overlay Plaque Assay. In M. R. J. Clokie & A. M. Kropinski (Eds.), Bacteriophages: Methods and Protocols, Volume 1: Isolation, Characterization, and Interactions (pp. 69–76). Humana Press. 10.1007/978-1-60327-164-6_7.

35. Bonilla, N., Rojas, M. I., Cruz, G. N. F., Hung, S.-H., Rohwer, F., & Barr, J. J. (2016). Phage on tap– a quick and efficient protocol for the preparation of bacteriophage laboratory stocks. PeerJ, 4, e2261. 10.7717/peerj.2261.

36. Boulanger, P. (2009). Purification of Bacteriophages and SDS-PAGE Analysis of Phage Structural Proteins from Ghost Particles. In M. R. J. Clokie & A. M. Kropinski (Eds.), Bacteriophages: Methods and Protocols, Volume 2 Molecular and Applied Aspects (pp. 227–238). Humana Press. 10.1007/978-1-60327-565-1_13.

37. Sasaki, M., Sitaraman, S. V., Babbin, B. A., Gerner-Smidt, P., Ribot, E. M., Garrett, N., Alpern, J. A., Akyildiz, A., Theiss, A. L., Nusrat, A., & Klapproth, J. M. (2007). Invasive Escherichia coli are a feature of Crohn’s disease. Laboratory investigation; a journal of technical methods and pathology, 87(10), 1042–1054. 10.1038/labinvest.3700661

38. Rousset M. (1986). The human colon carcinoma cell lines HT-29 and Caco-2: two in vitro models for the study of intestinal differentiation. Biochimie, 68(9), 1035–1040. 10.1016/s0300-9084(86)80177-8

39. Verhoeckx, K., Cotter, P., López-Expósito, I., Kleiveland, C., Lea, T., Mackie, A., Requena, T., Swiatecka, D., & Wichers, H. (Eds.). (2015). The Impact of Food Bioactives on Health: in vitro and ex vivo models. Springer.

40. Kucharzik, T., Hudson, J. T., Lügering, A., Abbas, J. A., Bettini, M., Lake, J. G., Evans, M. E., Ziegler, T. R., Merlin, D., Madara, J. L., & Williams, I. R. (2005). Acute induction of human IL-8 production by intestinal epithelium triggers neutrophil infiltration without mucosal injury. Gut, 54(11), 1565–1572. 10.1136/gut.2004.061168. .

41. Savidge, T. C., Newman, P. G., Pan, W.-H., Weng, M.-Q., Shi, H. N., McCormick, B. A., Quaroni, A., & Walker, W. A. (2006). Lipopolysaccharide-Induced Human Enterocyte Tolerance to Cytokine-Mediated Interleukin-8 Production May Occur Independently of TLR-4/MD-2 Signaling. Pediatric Research, 59(1), 89–95. 10.1203/01.pdr.0000195101.74184.e3.

42. Suzuki, M., Hisamatsu, T., & Podolsky, D. K. (2003). Gamma Interferon Augments the Intracellular Pathway for Lipopolysaccharide (LPS) Recognition in Human Intestinal Epithelial Cells through Coordinated Up-Regulation of LPS Uptake and Expression of the Intracellular Toll-Like Receptor 4- MD-2 Complex. Infection and Immunity, 71(6), 3503–3511. 10.1128/iai.71.6.3503-3511.2003

43. Gross, V., Andus, T., Daig, R., Aschenbrenner, E., Schölmerich, J., & Falk, W. (1995). Regulation of interleukin-8 production in a human colon epithelial cell line (HT-29). Gastroenterology, 108(3), 653–661. 10.1016/0016-5085(95)90436-0

44. Bille, E., Meyer, J., Jamet, A., Euphrasie, D., Barnier, J.-P., Brissac, T., Larsen, A., Pelissier, P., & Nassif, X. (2017). A virulence-associated filamentous bacteriophage of Neisseria meningitidis increases host-cell colonisation. PLOS Pathogens, 13(7), e1006495. 10.1371/journal.ppat.1006495

45. Nighot, M., Rawat, M., Al-Sadi, R., Castillo, E. F., Nighot, P., & Ma, T. Y. (2019). Lipopolysaccharide-Induced Increase in Intestinal Permeability Is Mediated by TAK-1 Activation of IKK and MLCK/MYLK Gene. The American journal of pathology, 189(4), 797–812. 10.1016/j.ajpath.2018.12.016

46. Stanifer, M. L., Guo, C., Doldan, P., & Boulant, S. (2020). Importance of Type I and III Interferons at Respiratory and Intestinal Barrier Surfaces. Frontiers in immunology, 11, 608645. 10.3389/fimmu.2020.608645.

47. Selsted, M. E., & Ouellette, A. J. (2005). Mammalian defensins in the antimicrobial immune response. Nature immunology, 6(6), 551–557. 10.1038/ni1206

48. Taniguchi, M., Okumura, R., Matsuzaki, T., Nakatani, A., Sakaki, K., Okamoto, S., Ishibashi, A., Tani, H., Horikiri, M., Kobayashi, N., Yoshikawa, H. Y., Motooka, D., Okuzaki, D., Nakamura, S., Kida, T., Kameyama, A., & Takeda, K. (2023). Sialylation shapes mucus architecture inhibiting bacterial invasion in the colon. Mucosal immunology, 16(5), 624–641. 10.1016/j.mucimm.2023.06.004

49. LeMessurier, K. S., Häcker, H., Chi, L., Tuomanen, E., & Redecke, V. (2013). Type I interferon protects against pneumococcal invasive disease by inhibiting bacterial transmigration across the lung. PLoS pathogens, 9(11), e1003727. 10.1371/journal.ppat.1003727

50. Watanabe, T., Asano, N., Fichtner-Feigl, S., Gorelick, P. L., Tsuji, Y., Matsumoto, Y., Chiba, T., Fuss, I. J., Kitani, A., & Strober, W. (2010). NOD1 contributes to mouse host defense against Helicobacter pylori via induction of type I IFN and activation of the ISGF3 signaling pathway. The Journal of clinical investigation, 120(5), 1645–1662. 10.1172/JCI39481

51. Barr, J. J., Auro, R., Furlan, M., Whiteson, K. L., Erb, M. L., Pogliano, J., Stotland, A., Wolkowicz, R., Cutting, A. S., Doran, K. S., Salamon, P., Youle, M., & Rohwer, F. (2013). Bacteriophage adhering to mucus provide a non–host-derived immunity. Proceedings of the National Academy of Sciences, 110(26), 10771–10776. 10.1073/pnas.1305923110

52. Zasloff, M. (2002). Antimicrobial peptides of multicellular organisms. Nature, 415(6870), Article 6870. 10.1038/415389a.

53. Turner, J., Cho, Y., Dinh, N.-N., Waring, A. J., & Lehrer, R. I. (1998). Activities of LL-37, a Cathelin-Associated Antimicrobial Peptide of Human Neutrophils. Antimicrobial Agents and Chemotherapy, 42(9), 2206–2214. 10.1128/aac.42.9.2206

54. Ganz, T. (2003). Defensins: Antimicrobial peptides of innate immunity. Nature Reviews Immunology, 3(9), Article 9. 10.1038/nri1180

55. O’Neil, D. A., Porter, E. M., Elewaut, D., Anderson, G. M., Eckmann, L., Ganz, T., & Kagnoff, M. F. (1999). Expression and Regulation of the Human β-Defensins hBD-1 and hBD-2 in Intestinal Epithelium1. The Journal of Immunology, 163(12), 6718–6724. 10.4049/jimmunol.163.12.6718.

56. Schauber, J., Svanholm, C., Termén, S., Iffland, K., Menzel, T., Scheppach, W., Melcher, R., Agerberth, B., Lührs, H., & Gudmundsson, G. H. (2003). Expression of the cathelicidin LL-37 is modulated by short chain fatty acids in colonocytes: Relevance of signalling pathways. Gut, 52(5), 735–741. 10.1136/gut.52.5.735.

57. Jiang, W., Sunkara, L. T., Zeng, X., Deng, Z., Myers, S. M., & Zhang, G. (2013). Differential regulation of human cathelicidin LL-37 by free fatty acids and their analogs. Peptides, 50, 129–138. 10.1016/j.peptides.2013.10.008.

58. Dalile, B., Van Oudenhove, L., Vervliet, B., & Verbeke, K. (2019). The role of short-chain fatty acids in microbiota–gut–brain communication. Nature Reviews Gastroenterology & Hepatology, 16(8), Article 8. 10.1038/s41575-019-0157-3.

59. Cummings, J. H., Pomare, E. W., Branch, W. J., Naylor, C. P., & Macfarlane, G. T. (1987). Short chain fatty acids in human large intestine, portal, hepatic and venous blood. Gut, 28(10), 1221–1227. 10.1136/gut.28.10.1221.

60. Choi, J., & Augenlicht, L. H. (2024). Intestinal stem cells: Guardians of homeostasis in health and aging amid environmental challenges. Experimental & Molecular Medicine, 56(3), 495–500. 10.1038/s12276-024-01179-1

61. Zamora, P. F., Reidy, T. G., Armbruster, C. R., Sun, M., Van Tyne, D., Turner, P. E., Koff, J. L., & Bomberger, J. M. (2024). Lytic bacteriophages induce the secretion of antiviral and proinflammatory cytokines from human respiratory epithelial cells. PLoS biology, 22(4), e3002566. 10.1371/journal.pbio.3002566

62. Miernikiewicz, P., Kłopot, A., Soluch, R., Szkuta, P., Kęska, W., Hodyra-Stefaniak, K., Konopka, A., Nowak, M., Lecion, D., Kaźmierczak, Z., Majewska, J., Harhala, M., Górski, A., & Dąbrowska, K. (2016). T4 Phage Tail Adhesin Gp12 Counteracts LPS-Induced Inflammation In Vivo. Frontiers in Microbiology, 7. 10.3389/fmicb.2016.01112

63. Zhang, L., Hou, X., Sun, L., He, T., Wei, R., Pang, M., & Wang, R. (2018). Staphylococcus aureus Bacteriophage Suppresses LPS-Induced Inflammation in MAC-T Bovine Mammary Epithelial Cells. Frontiers in Microbiology, 9. 10.3389/fmicb.2018.01614

64. Washizaki, A., Yonesaki, T., & Otsuka, Y. (2016). Characterization of the interactions between Escherichia coli receptors, LPS and OmpC, and bacteriophage T4 long tail fibers. MicrobiologyOpen, *5*(6), 1003–1015. 10.1002/mbo3.384

65. Feige, U., & Stirm, S. (1976). On the structure of the Escherichia coli C cell wall lipopolysaccharide core and on its øX174 receptor region. Biochemical and Biophysical Research Communications, 71, 566–573.

66. Bruno, M. E. C., Rogier, E. W., Arsenescu, R. I., Flomenhoft, D. R., Kurkjian, C. J., Ellis, G. I., & Kaetzel, C. S. (2015). Correlation of Biomarker Expression in Colonic Mucosa with Disease Phenotype in Crohn’s Disease and Ulcerative Colitis. Digestive Diseases and Sciences, 60(10), 2976– 2984. 10.1007/s10620-015-3700-2.

67. Lin, X., Li, J., Zhao, Q., Feng, J., Gao, Q., & Nie, J. (2018). WGCNA Reveals Key Roles of IL8 and MMP-9 in Progression of Involvement Area in Colon of Patients with Ulcerative Colitis. Current Medical Science, 38(2), 252–258. 10.1007/s11596-018-1873-6.

68. Korolkova, O. Y., Myers, J. N., Pellom, S. T., Wang, L., & M’Koma, A. E. (2015). Characterization of Serum Cytokine Profile in Predominantly Colonic Inflammatory Bowel Disease to Delineate Ulcerative and Crohn’s Colitides. Clinical Medicine Insights. Gastroenterology, 8, 29–44. 10.4137/CGast.S20612.

69. Tenaillon, O., Skurnik, D., Picard, B., & Denamur, E. (2010). The population genetics of commensal Escherichia coli. Nature Reviews Microbiology, 8(3), Article 3. 10.1038/nrmicro2298

70. Wells, C. L., Jechorek, R. P., Olmsted, S. B., & Erlandsen, S. L. (1993). Effect of LPS on epithelial integrity and bacterial uptake in the polarized human enterocyte-like cell line Caco-2. Circulatory shock, 40(4), 276–288.

71. Wells, CL, Jechorek RP, Olmsted SB, and Erlandsen SL. (1994). Bacterial Translocation In Cultured Enterocytes: Magnitude, Specificity, And Electron Microscopic Observations Of Endocytosis. Shock 1(6):p 443–451.

72. Olmsted, S. B., Dunny, G. M., Erlandsen, S. L., & Wells, C. L. (1994). A plasmid-encoded surface protein on Enterococcus faecalis augments its internalization by cultured intestinal epithelial cells. The Journal of infectious diseases, 170(6), 1549–1556. 10.1093/infdis/170.6.1549

73. Popescu, M. C., Haddock, N. L., Burgener, E. B., Rojas-Hernandez, L. S., Kaber, G., Hargil, A., Bollyky, P. L., & Milla, C. E. (2024). The Inovirus Pf4 Triggers Antiviral Responses and Disrupts the Proliferation of Airway Basal Epithelial Cells. Viruses, 16(1), Article 1. 10.3390/v16010165

74. Ning, S., Pagano, J. S., & Barber, G. N. (2011). IRF7: activation, regulation, modification and function. Genes and immunity, 12(6), 399–414. 10.1038/gene.2011.21

75. Liu, Y., Yu, Z., Zhu, L., Ma, S., Luo, Y., Liang, H., Liu, Q., Chen, J., Guli, S., & Chen, X. (2023). Orchestration of MUC2 — The key regulatory target of gut barrier and homeostasis: A review. International Journal of Biological Macromolecules, 236, 123862. 10.1016/j.ijbiomac.2023.123862

76. Chin, W. H., Kett, C., Cooper, O., Müseler, D., Zhang, Y., Bamert, R. S., Patwa, R., Woods, L. C., Devendran, C., Korneev, D., Tiralongo, J., Lithgow, T., McDonald, M. J., Neild, A., & Barr, J. J. (2022). Bacteriophages evolve enhanced persistence to a mucosal surface. Proceedings of the National Academy of Sciences of the United States of America, 119(27), e2116197119. 10.1073/pnas.2116197119

77. Tian, Y., Wu, M., Liu, X., Liu, Z., Zhou, Q., Niu, Z., & Huang, Y. (2015). Probing the Endocytic Pathways of the Filamentous Bacteriophage in Live Cells Using Ratiometric pH Fluorescent Indicator. Advanced Healthcare Materials, 4(3), 413–419. 10.1002/adhm.201400508.

78. Lehti, T. A., Pajunen, M. I., Skog, M. S., & Finne, J. (2017). Internalization of a polysialic acid- binding Escherichia coli bacteriophage into eukaryotic neuroblastoma cells. Nature communications, 8(1), 1915. 10.1038/s41467-017-02057-3

79. Hsia, R., Ohayon, H., Gounon, P., Dautry-Varsat, A., & Bavoil, P. M. (2000). Phage infection of the obligate intracellular bacterium, Chlamydia psittaci strain guinea pig inclusion conjunctivitis. Microbes and infection, 2(7), 761–772. 10.1016/s1286-4579(00)90356-3

